# Estradiol and Flutamide Effects on the Song System of Developing Male Zebra Finches

**DOI:** 10.1101/2024.03.11.584474

**Authors:** William Grisham, Mary Ellen McCormick

## Abstract

Estradiol (E_2_) masculinizes the developing song system of female zebra finches (Taeniopygia castanotis) if it is administered in early life, but its effect is blocked with the co-administration of an antiandrogen (Flutamide). The effects of E_2_ on the developing male song system are not uniform and reports of Flutamide administration in developing male zebra finches differ in their findings. Therefore. this study was conducted to further explore the effects of administering E_2_ alone, Flutamide (Flut) alone, or the two in combination during early post-hatch development. Brains and testes were examined after day 100. The results showed definite demasculinizing effects of early E_2_ on the song nucleus HVC (proper name)—its volume and neuron number were markedly reduced. Nonetheless, early E_2_ hypermasculinized HVC neuronal size. Flut slightly hypermasculinized RA volume (Robust nucleus of the Arcopallium), which replicates one prior study but the absence of additional effects is at odds with others. Early E2 resulted in markedly reducing testes size, which is likely to be a consequence of hijacking endogenous endocrine feedback mechanisms. Arguments are put forward suggesting 1) early E2 action on HVC could be an anachronistic consequence of actions on the genotype of developing male versus females or 2) a disruption of endocrine mechanisms inducing inappropriate hormonal states during development. These possibilities are not mutually exclusive.

## Introduction

Song behavior in zebra finches (Taeniopygia castanotis) is sexually dimorphic; males sing and females do not. This sex difference in behavior is reflected by dramatic sex differences in its neural underpinnings. Males have larger song nuclei such as HVC, RA—robust nucleus of the arcopallium, and Area X (Grisham & Arnold, 1995), with more neurons in HVC and RA (Nordeen & Nordeen, 1988; Schlinger & Arnold, 1991), and larger neurons in HVC, lMAN—Lateral Magnocellular nucleus of the Anterior Nidopallium, and RA (Nottebohm & Arnold, 1976; Grisham & Arnold, 1995; Arnold & Saltiel, 1979; Nordeen et al., 1992).

The female’s song system can be masculinized by administering estradiol (E_2_) during development (Adkins-Regan et al., 1994; Gurney & Konishi, 1980; Gurney, 1981, 1982; Jacobs, Grisham & Arnold, 1995; Simpson & Vicario, 1991). The influence of early E_2_ on song system development in males is less clear, however.

Two studies report that E_2_ has no effect on the song system of developing male zebra finches either when administered at hatching (Gurney & Konishi, 1980) or *in ovo* (Wade et al., 1997). Nonetheless, Tang and Wade (2009) found early E_2_ in males increased Area X volume when examined at day 25 posthatch but had no impact on HVC. Conversely, when males were treated for the first 25 days posthatch with estradiol benzoate (EB) and examined at day 25, the volume of Area X and HVC were demasculinized (Mathews & Arnold, 1991), but not when males were treated until day 20 then examined at day 60 when early EB treatment hypermasculinized male neuronal sizes in lMAN and HVC (Mathews & Arnold, 1991).

Mixed effects of androgenic manipulations in development also have been reported. Blocking androgen action with Flutamide (Flut) at hatching hypermasculinized RA volume and the number of its neurons (Schlinger & Arnold, 1991). But, life-long exposure to Flut demasculinized RA neuron number and slightly decreased HVC volume (Grisham et al., 2007). Also, when it was administered at day 20 to castrates, it demasculinized both lMAN and Area X but did not influence other song regions (Bottjer & Hewer, 1992). Blocking synthesis of an androgen, dihydrotestosterone, at hatching reduced the number and density of RA neurons (Grisham et al., 1997).

Administering Flut at hatching along with E_2_ blocked the usual E_2_ induced masculization of female zebra finch song system (Grisham et al., 2002). Given the mixed results of both estrogen and androgenic manipulations of males in early life, we decided to re-examine the effects of these treatments as well as their combination. We assessed the effects of both E_2_ and Flut alone and in combination on hatchling males, particularly since we had found such dramatic effects on the combined treatment in hatchling females.

## Methods

All aspects of this study were approved by the Animal Research Committee of the University of California, Los Angeles. Further, the ARRIVE guidelines 2.0 were followed.

Estradiol (E_2_) and flutamide (Flut) were administered in a 2×2 design. Males were implanted within 1-3 days of hatching under the skin of the breast with E_2_ pellets, Flut pellets, both, or neither. The E_2_ implants were made by mixing finely ground E_2_ with Medical grade Silastic glue in a ratio of 1:6 and extruding this mixture into polyethylene tubing (Clay Adams no. 7411, i.d. 0.58 mm, o.d. 0.965 mm). These implants were cut into 2 mm sections, weighed, and their average dose calculated to be 83 µg (batches of pellets ranged from 73 µg to 87 µg E_2_). The Flut implants were made in a manner described by Schlinger and Arnold (1991). Briefly, Flut was mixed with Silastic glue and then extruded through a 1cc syringe without a needle. The resulting strips were left to cure overnight, quartered, weighed, and cut into lengths that would result in a dose of 200 µg. Each bird received two pellets (E_2 +_ blank, E_2_ + Flut, Flut + blank, or two blanks) except for unimplanted controls.

All dependent measures were made blind to the birds’ treatment group. At sacrifice (101-136 days of age), the birds were deeply anesthetized and then perfused with 0.75% saline followed by 10% formalin in saline. Birds were only included if the pellets could be found at sacrifice. (Birds in the blank+blank group were included regardless.) Even after having been implanted for 101 days or more, the E_2_ pellets all appeared to contain E_2_ crystals (they had white opacities). The Flut pellets were clear and appeared depleted. The testes were dissected and stored in 10% formalin until they were weighed; at which time they were cleaned of extraneous tissue, dabbed dry, and weighed to the nearest 0.1 mg (if testes weighed less than .1 mg, they were assigned a 0.1 mg weight). Testes weights were taken from 26 adult males (n = 5 E_2_ treated; n = 4 E_2_+Flut treated; n=8 Flut treated; and 9 untreated/blanks) and both testes were averaged for each individual.

The brains were dissected, stored in 10% formalin, and frozen-sectioned at 40 µm in the coronal plane. Brain measures were taken from 28 individuals (n = 5 E_2_-treated; n = 4 E_2_+Flut treated; n = 8 Flut treated; n = 11 untreated/blanks), but due to histological problems HVC data was n = 26. Data and records can be accessed via UCLA Dataverse at https://doi.org/10.25346/S6/VNSKRP.

The brains were examined with a light microscope connected to a computer via a videocamera. Cross-sectional areas of song system nuclei and their individual neurons were measured using NIH Image (http://rsb.info.nih.gov/nih-image/).

The volumes of Area X, lMAN, HVC, and RA were calculated by means of the cylindrical method: tracing the cross-sectional areas on every third section magnified at 6.25X, adding these areas, and multiplying by the sampling interval (120 µm). The cross-sectional volumes of each song system nucleus were averaged across hemispheres for each animal unless one hemisphere was damaged.

The area of individual neurons was measured at 800X. Twenty-five neurons were sampled through the rostral-caudal extent of the nucleus in each hemisphere for a total of fifty neurons in each song control nucleus of each animal. Neurons were distinguished from glia by their dark staining, ample cytoplasm, and nuclei containing only one or two nucleoli. Glia were distinguished by light staining, little cytoplasm, often with several nucleoli in each nucleus. The number of neurons in HVC was determined by counting the number of nucleoli in twenty-five frames (each 45,350 μm^3^) that were sampled throughout its rostral-caudal extent in both hemispheres. Counting a small profile like nucleoli produces counts as reliable as using an optical dissector (Tramontin et al., 1998). The average density of neurons was calculated for each animal and multiplied by the mean volume of the given song system nucleus.

Dependent variables were analyzed via JASP https://jasp-stats.org/ and using a 2×2 ANOVA: estradiol treatment vs. none as one factor and Flut treatment vs. none as the other factor. Testes weights were analyzed by an ANOVA with 2×2 between (E_2_ vs. none and flut vs. none) and one within variable (side).

## Results

HVC volume was significantly demasculinized (reduced) by E_2_ treatment, F(1,22) = 8.38, <u>p</u> < 0.01, η^2^ = 0.272 (Figure 1A), as was the number of HVC neurons, F(1,22) = 4.51, <u>p</u> < 0.05, η^2^ = 0.168 (Figure 1B). Despite E_2_’s demasculinizing effect on the volume of HVC and on the number of its neurons, it hypermasculinized the size of HVC neurons, F(1,24) = 6.00, <u>p</u> < 0.05, η^2^ = 0.195 (Figure 1C). There was a weak effect/strong trend for Flut hypermasculinizing RA volume, F(1,24) = 4.133, p = 0.053, η^2^ = 0.131 (Figure 1D). No other brain measures were affected by either E_2_ or Flut alone or in combination (all p values > 0.09).

**Figure 1.**
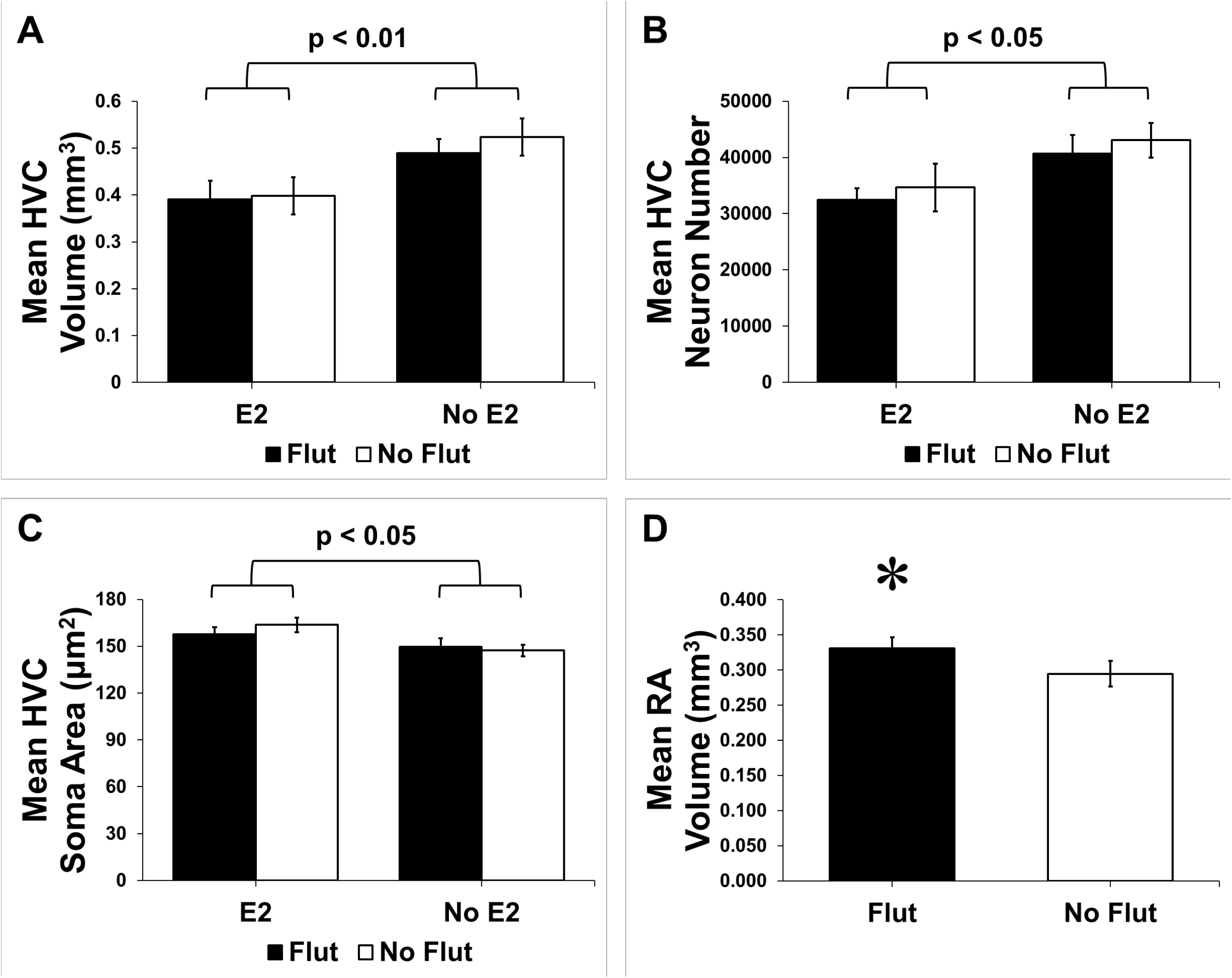
A) Mean HVC volume and B) number of HVC neurons as a function of treatment group. Males receiving E_2_ or E_2_ +Flut were demasculized. C) Mean HVC neuron size—males in E_2_-treated groups had significantly larger neurons. D) RA volume--males receiving Flut were hypermasculinized * indicates a strong trend p < 0.06. Error bars = SEM.

Testes weight was markedly reduced by E_2_ treatment, F(1,69) = 74.63, <u>p</u> < 0.0001, η ^2^ = 0.475 (Figure 2A). There was no effect of Flut alone or an E_2_ x Flut interaction on testes size, all <u>p</u>s > 0.50 (Figure 2A). Both HVC volume and the number of HVC neurons significantly correlated with mean testicular weight, r(22) = .576, <u>p</u> < 0.01 and r(22) = .487, <u>p</u> < 0.02, respectively (Figure 2B & 2C).

**Figure 2.**
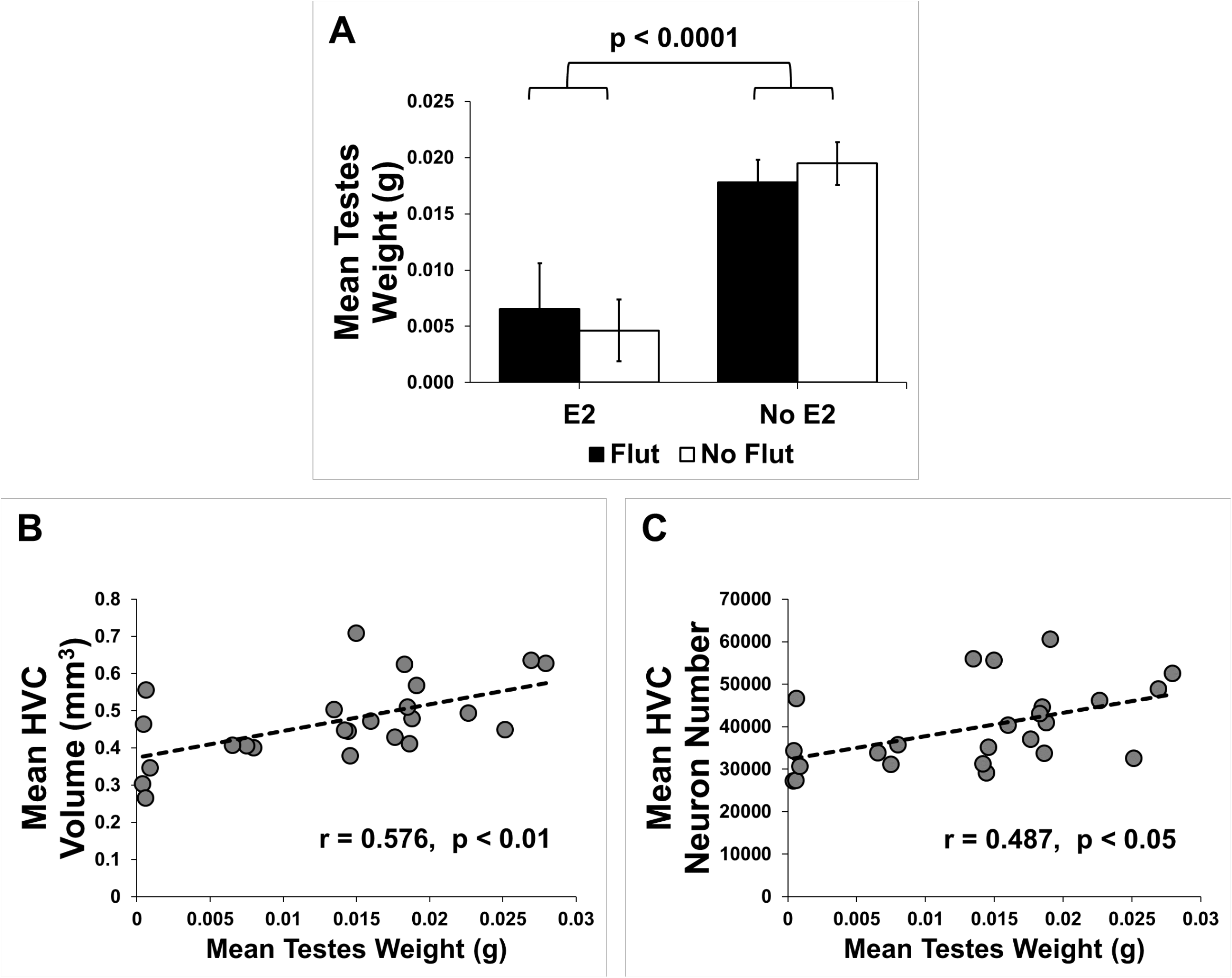
A) Mean testes weight as a function of treatment--E_2_ treated birds had markedly reduced testes weights. Error bars = SEM. B) Scatterplot of HVC volume and C) HVC neuron numbers as a function of testes size.

## Discussion

Early administration of E_2_ to male hatchlings dramatically reduced HVC volume and the number of its neurons when examined in adulthood (Fig. 1A, 1B), which contrasts sharply with the masculinizing effects of early E_2_ in females (Adkins-Regan et al., 1994; Grisham & Arnold, 1995; Gurney & Konishi, 1980; Gurney, 1981, 1982; Jacobs et al., 1995; Simpson & Vicario, 1991). Early E_2_ administration to males could 1) differentially affect the male and female genome, possibly triggering temporally inappropriate genetic cascades, and/or 2) interfer with hormonal feedback mechanisms and gonadal function.

There is evidence of early E_2_ differentially altering BDNF gene expression in male but not female HVC (Dittrich et al., 1999). This result suggests that early E_2_ disrupts the temporal sequence of male gene expression, which could alter the adult phenotype. Early E_2_ administration also has opposite effects on gene expression in females versus males; the number of HVC neurons expressing 17β-hydroxysteroid dehydrogenase mRNA decreased in males but increased in females when birds were examined at day 25 (Thompson et al., 2011). Early E_2_ also alters genes in female Area X toward the masculine pattern whereas it induces more female-like patterns in male lMAN and RA (Choe et al., 2021), the latter of which is a target of HVC projections (Benezra et al., 2018). Thus, early E_2_ could have its action on male HVC via its axonal projections to RA.

Temporal differences in early E_2_ administration effects may explain some discrepancies in the literature. Notably, E_2_ at hatching decreases the number of BrdU-labeled cells added to HVC in males at puberty whereas it increases the number of labeled cells in females (Tang and Wade, 2009). Choe et al. (2021) did not detect any effect of early E_2_ on male HVC volume, but we did. Nonetheless, several methodological differences could explain this discrepancy: different dose regimens; different ways of defining HVC, and very different time points when the song systems were measured (day 30, vs. > 100 days of age). HVC neurons are added well after day 30 (Walton et al., 2012; Diez et al., 2021), which could also explain the discrepancy between the data of Choe et al. (2021) and ours. In agreement with our findings, however, Choe et al. (2021) also found early E_2_ increased HVC cell size in males.

Hormonal mechanisms could have played a role on the brain and gonadal development in the present study. The dramatic decrease in testes size induced by early E_2_ administration (Fig. 2A) suggests that there are estrogen-mediated endocrine feedback mechanisms in zebra finches. This finding is paralleled in other studies on zebra finches (Mathews & Arnold, 1991), and other bird species (Casto & Ball, 1996; Lorenz, 1954; Soma et al., 2000). So, our early E_2_ exposure could have permanently altered hormonal feedback mechanisms resulting in small testicular size.

Notably, HVC volume and the number of its neurons were significantly correlated with testes size (Fig 2B, 2C). The testes in our early E_2_-treated birds were more like those of babies rather than adults suggesting that they never matured. The best estimate of the onset of puberty in zebra finches is about Day 60-70 posthatch (Bölting & von Englehardt, 2017; Pröve, 1983) and we did not examine the birds until after Day 100.

HVC neurons are still being added up until Day 70 (Diez et al., 2021) and even into adulthood (Walton et al., 2012). Testosterone increases the recruitment and/or survival of HVC neurons in adult female canaries (Rasika et al., 1994), an effect that is dependent upon the androgenic rather than estrogenic metabolites of testosterone (Fusani et al., 2002). If the markedly reduced testes size in our birds resulted in reduced testicular secretions not only before but also after puberty, then demasculinizing effects could be a consequence.

The number of androgen target cells in HVC normally increases during adolescence in male zebra finches (Bottjer, 1987; Tang & Wade, 2010). Early E_2_ treatment decreases androgen receptor levels in male but not female HVC (Thompson et al., 2011), so there may have been not only low androgen levels, but also fewer receptors upon which to act.

Our early antiandrogen (Flut) treatment showed a strong trend toward hypermasculinizing RA volume (Figure 1D) similar to Schlinger and Arnold (1991)—we used the same protocol as they did. We did not replicate the Flut effects of Bottjer and Hewer (1992) or Grisham et al. (2007) on the song system or on testes size (Fig. 2A), but the protocol of these other studies was quite different in timing of Flut administration. Our Flut pellets in this experiment were likely to be depleted before the birds were examined, which could explain the difference in outcomes.

In conclusion, the alteration of the male song system by early E_2_ or Flut may be explained either by E_2_ regulating genes in abberant fashions and/or by the altering steroid hormone profile of the developing males. The latter would probably result from the apparent lack of testes development, which would lead to a crucial lack of androgens impacting HVC development. These two explanations are not mutually exclusive, and in fact may be intertwined. Consideration of timing and duration of treatments as well as impacts on endocrine systems in development could unconfound the discrepancies in the literature.

## Acknowledgements

Thanks to Dr. Juli Wade and Dr. Arthur P. Arnold who provided valuable feedback on an earlier version of this manuscript. Further thanks to Dr. Art Arnold for allowing us to use his facilities. Thanks to Natalie Schottler who drew the figures and made the final edits and Kay Yang-Stayner and Janet Lee who helped with the execution of the study. This research did not receive any specific grant from funding agencies in the public, commercial, or not-for-profit sectors.

## Notes

https://doi.org/10.25346/S6/VNSKRP

### Competing Interest Statement

The authors have declared no competing interest.

### Summary of Updates

Better description of the question and discussion of results.

